# Proteolysis targeting chimera extracellular vesicles for therapeutic development treating triple negative breast cancer

**DOI:** 10.1101/2024.08.25.609564

**Authors:** Nina Erwin, Umasankar De, Yufeng Xiao, Lei Wang, Chandra Maharjan, Xiaoshu Pan, Nikee Awasthee, Guangrong Zheng, Daiqing Liao, Weizhou Zhang, Mei He

## Abstract

Proteolysis targeting chimeras (PROTACs) are an emerging targeted cancer therapy approach, but wide-spread clinical use of PROTAC is limited due to poor cell targeting and penetration, and instability in vivo. To overcome such issues and enhance the in vivo efficacy of PROTAC drugs, microfluidic droplet-based electroporation (µDES) was developed as a novel extracellular vesicle (EVs) transfection system, which enables the high-efficient PROTAC loading and effective delivery in vivo. Our previously developed YX968 PROTAC drug had shown the selectively degradation of HDAC3 and 8, which effectively suppresses the growth of breast tumor cell lines, including MDA-MB-231 triple negative breast cancer (TNBC) line, via dual degradation without provoking a global histone hyperacetylation. In this study, we demonstrated that µDES-based PROTAC loading in EVs significantly enhanced therapeutic function of PROTAC drug in vivo in the TNBC breast tumor mouse model. NSG mice with pre-established MDA-MB-231 tumors and treated with intraperitoneal injection of EVs for tumor inhibition study, which showed significantly higher HDAC 3 and 8 degradation efficiency and tumor inhibition than PROTAC only group. The liver, spleen, kidney, lung, heart, and brain were collected for safety testing, which exhibited improved toxicity. The EV delivery of PROTAC drug enhances drug stability and bioavailability in vivo, transportability, and drug targeting ability, which fills an important gap in current development of PROTAC therapeutic functionality in vivo and clinical translation. This novel EV-based drug transfection and delivery strategy could be applicable to various therapeutics for enhancing in vivo delivery, efficacy, and safety.

## Introduction

Breast cancer is a significant global health concern, due to the large number of reported incidences worldwide, as the top leading cause of cancer mortality in women^1-4^. Breast tumors are incredibly heterogenous with diverse molecular features, different sensitivities to therapies, and various prognoses, which requires advanced treatment strategies against breast cancer^1-3, 5^. The triple negative breast cancer (TNBC) subtype is especially challenging for therapeutic development, due to the aggressiveness of high proliferation rates and early metastasis, and substantial proneness to relapse^2, 3, 6^. Standard targeted breast cancer treatment regimens are often failed in treating TNBC ^2, 3, 6, 7^ with poor therapeutic responses.^2, 3, 6, 7^. It has been found that class I histone deacetylases, including HDAC2, 3, and 8, are significantly overexpressed in breast tumor tissue relative to normal breast tissue^8^. The well-established HDAC mediated mechanism for cancer drug development has led to the use of HDAC inhibitors (HDACi). However, most HDACis are pan-histone deacetylase inhibitors, which often nonspecifically lead to global hyperacetylation, contributing to off target toxicity effects^8-12^ and ineffectiveness in targeting solid tumors^11, 13, 14^ in breast cancer. Recently, proteolysis targeting chimera (PROTAC) has been emerging as the new therapeutic drug targeting the catalysis of histone deacetylases (HDAC), which enables the degradation of HDACs for activating tumor suppressor genes and resulting in the antitumor effects^4, 15^. Regardless of this promising development of PROTAC as a breast cancer therapeutic drug, significant drawbacks remain limiting the clinical use. PROTAC molecules have a relatively high molecular weight and large exposed polar surface area^16, 17^. Such unideal physiochemical and pharmacokinetic properties of PROTAC could lead to poor absorption and cell penetration, limiting the bioavailability in tumor tissues (**Fig. 1**)^17^. Herein, we developed extracellular vesicle enabled PROTAC delivery which improves in vivo efficacy of PROTAC therapeutic functional for broader clinical anti-cancer applications, owning to their high biocompatibility, long circulation times, and inherent targeting and tissue penetration (**Fig. 1**) ^18-21^.

**Figure 1.**
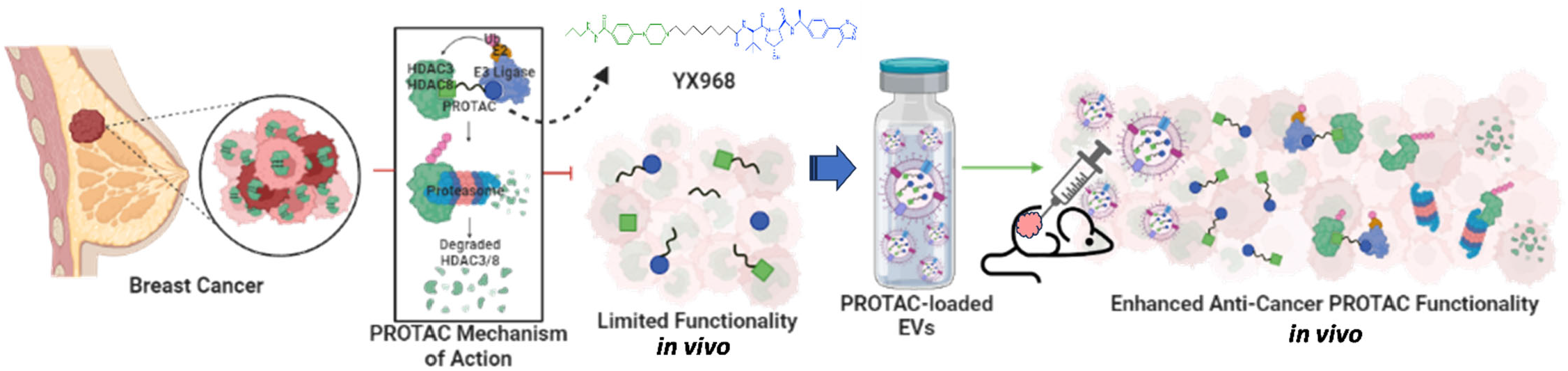
Breast cancer with overexpressed HDAC3 and HDAC8 proteins can be treated with YX968 PROTAC-loaded EVs for the efficient in vivo delivery, leading to proteasomal degradation and anti-tumor effects.

We have pioneered a development of isozyme-specific PROTAC YX968, which displayed dual degradation selectively on both HDAC3 and 8, and effectively suppressed the growth of various breast cancer cell lines, particularly the MDA-MB-231 TNBC cell line^22^. However, the in vivo tumor inhibition study is not successful. We suspect it may be due to poor absorption and cell penetration, limiting the bioavailability in the tumor site in vivo. In order to overcome such challenges from in vivo delivery, in this study, we introduced a novel microfluidic droplet-based electroporation (µDES) system for EV transfection, which enables the high-efficient PROTAC loading into EVs while retaining high integral properties of EVs. Droplet-based continuous-flow microfluidic electroporation also offers high throughput, low-voltage for retaining EV quality and integrity, and low-cost production. We observed that µDES produced PROTAC EVs significantly enhanced PROTAC therapeutic function in vivo in the TNBC breast tumor mouse model. NSG mice with pre-established MDA-MB-231 tumors were treated with intraperitoneal injection of XY968 PROTAC EVs for tumor inhibition study, which exhibited significantly higher HDAC 3 and 8 degradation efficiency and tumor inhibition than XY968 PROTAC only group. The liver, spleen, kidney, lung, heart, and brain were collected for safety testing, which displayed improved toxicity profile. The EV delivery of PROTAC drug enhances drug stability and bioavailability in vivo which will fill an important gap in current development of PROTAC therapeutic functionality in vivo and clinical translation. This novel EV-based drug transfection and delivery strategy will be appliable broadly to various therapeutics for enhancing in vivo delivery, efficacy, and safety.

## Results

### Characterization of transfection methods for producing high quality PROTAC EVs

Various methods can be employed to load PROTAC molecules into EVs, including passive incubation, which is a simple diffusion-based method and often leads to low drug incorporation and loading stability^18, 20, 23^. Alternatively, active loading strategies, including chemically induced cationic transfection, can be employed^24^. In this method, commercial transfection reagents, like lipofectamine, can merge with the EV membrane through electrostatic interactions^25^. However, various EV characteristics are altered as an effect, with various changes to EV size and surface charge properties^21, 23, 24^, as well as potential toxicity effects from lipofectamine^26^. Electroporation has been shown to be a successful loading technique, using electrical fields to induce temporary pores formation in the EV membrane and improve drug transfection^18, 20, 21, 25-28^. We investigated the influences of various transfection methods on EVs’ characteristics following PROTAC loading, which is critical to develop the optimal in vivo delivery and tissue targeting profile (**Fig.2A**). The potential alterations in quality of EVs and their surface properties could affect EVs’ efficacy as a drug delivery vehicle in vivo. As shown in **Fig.2B**, no noticeable changes were observed in terms of EV size following the transfection techniques. However, variability was present for the EVs regarding their particle concentration (**Fig. 2C**) and zeta potential (**Fig. 2D**). Particularly for lipofectamine transfected EVs, the marked changes showed a decline in particle concentration and zeta potential, compared to the unloaded native EVs. The lipofection induced alterations were further proved through morphological assessments using Transmission Electron Microscopy (TEM) imaging, showing rough surface properties and degradation. These alterations could be due to the membrane damage often associated with cationic liposome interactions with negatively charged EVs^29^. EVs transfected through incubation and traditional electroporation showed that EVs maintained similar size, shape, and nanostructure as the unloaded native EVs, which provides insights into their potential functional applications.

**Figure 2.**
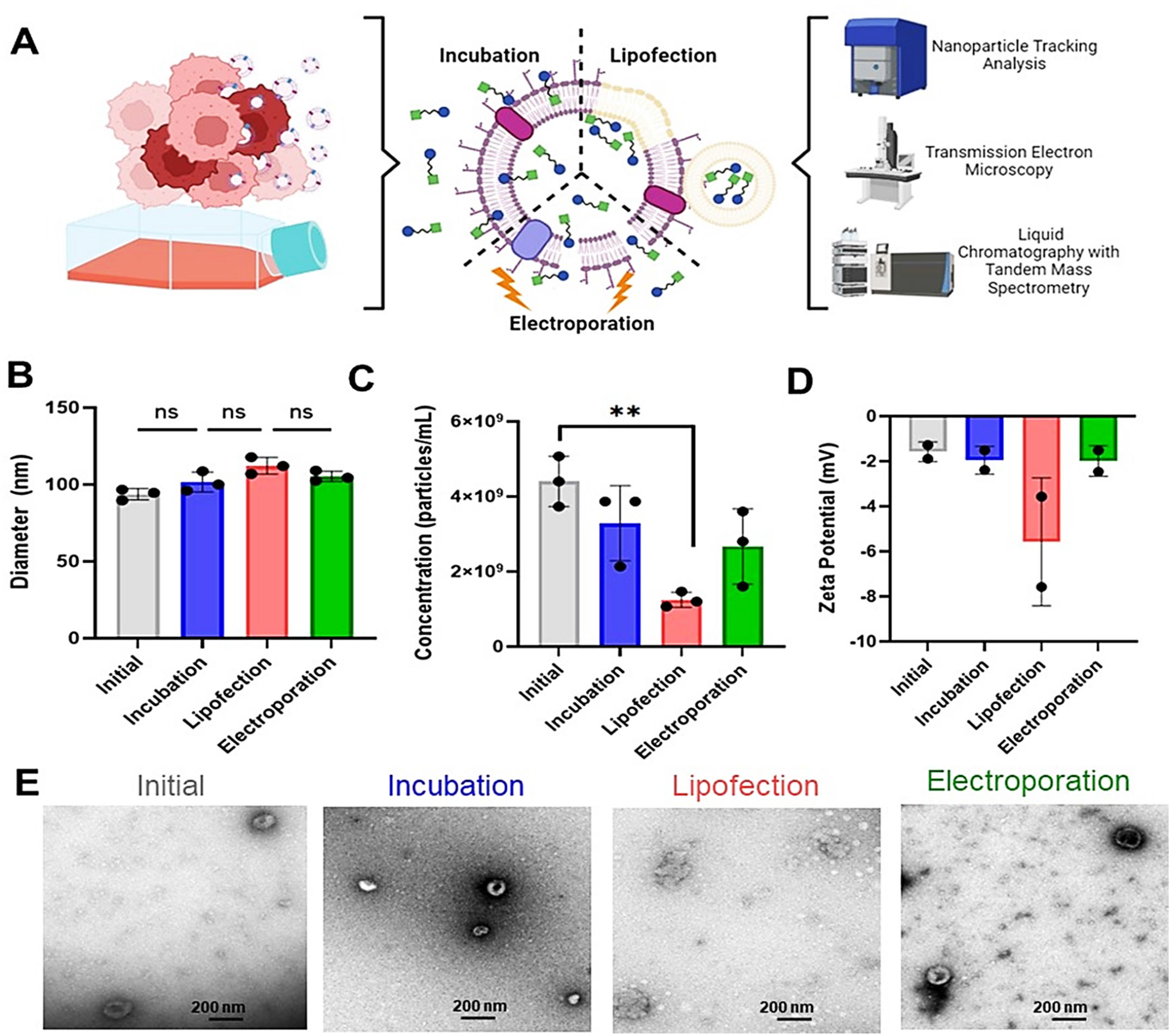
A) Workflow depicting breast cancer derived EV isolation for PROTAC loading using conventional transfection methods and subsequent characterization to assess the effect of loading on EV B) size, C) concentration, D) zeta potential, and E) morphology.

While conventional cuvette electroporation has promising potential as a transfection technique for EVs with much higher loading capacity, the method is limited by low-throughput rates for clinical translation. The high voltage conditions in kilowatts often leads to EV aggregation and toxicity^18, 20, 21, 23, 25, 27, 28^. In efforts to improve electroporation strategies for enhancing PROTAC cargo loading and transforming drugs to clinical therapeutics, we introduced a novel continuous-flow microfluidic droplet-based electroporation (μDES). In this method, hydrodynamic forces and interfacial tension at a junction between phases will lead to the formation of thousands of buffer in oil droplets per second, and then flow through electrodes surrounded microfluidic channel side for electroporation (**Fig. 3A**)^30, 31^. Droplet-based continuous-flow microfluidic electroporation offers high throughput, low-voltage for retaining EV quality and integrity, low-cost production, and simple operation. The system is also highly customizable, as flow rates, pressures, and voltages can all be adjusted for rapid, precise, and highly reproducible droplet formation and EV transfection (**Fig. 3B**)^30, 32^. Additionally, due to the large surface-area-to-volume ratios of the droplets, mass transport is highly efficient, allowing for enhancing transfection efficiency and reducing the membrane damage or alteration^27, 28, 30, 31^. Following the implementation of this novel transfection technique, EVs’ integral properties were characterized in Figure 3, compared to conventional electroporation and the native EVs as the control group. No noticeable changes were observed in terms of EV size (**Fig. 3C**), concentration (**Fig. 3D**), zeta potential (**Fig. 3E**), or morphology (**Fig. 3F**) following μDES based PROTAC loading.

**Figure 3.**
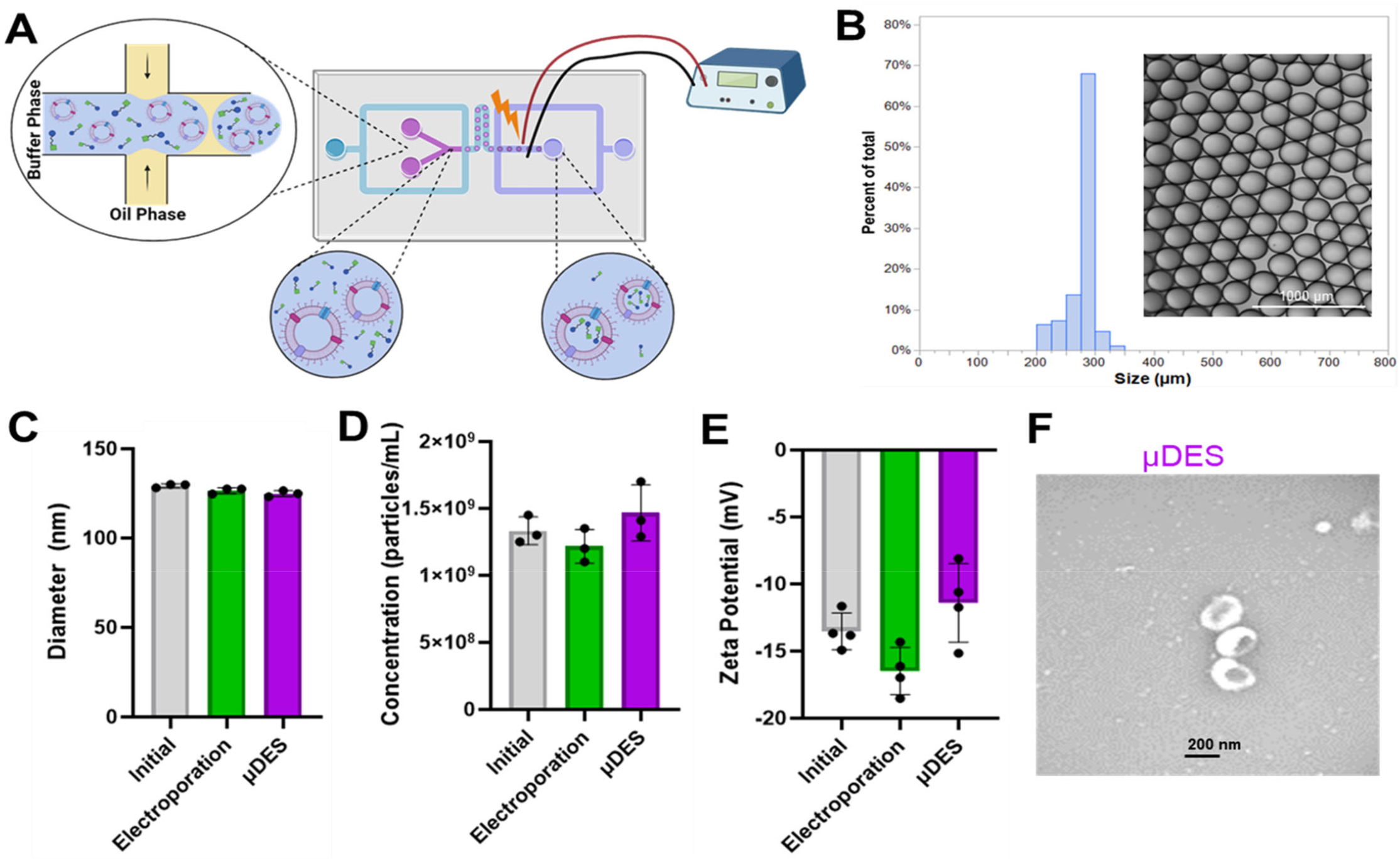
A) Schematic of the novel microfluidic droplet-based electroporation (μDES) device for PROTAC loading into EVs. B) Uniform size and morphology of the droplets produced following μDES. Insert is the bright field image of produced droplets. The transfected PROTAC EVs were characterized in terms of C) size, D) concentration, E) zeta potential, and F) morphology, compared to conventional electroporation EVs and native EVs as the control group.

For optimal functionality as drug delivery vehicles in vivo, the transfection efficiency of PROTAC into EVs, as well as the stability of the loaded PROTAC EVs under different storage conditions, were characterized. The transfection efficiency was measured by HPLC-MS on the loaded drug fraction after the removal of unincorporated PROTAC and normalized to the molar concentration of PROTAC input in the transfection solution. EVs that did not undergo any PROTAC loading were used as a control. Each of the loading methods demonstrated distinct loading efficiencies, with chemical transfection resulting in the highest loading rates of the investigated methods (**Fig. 4A**). The high loading efficiency following chemical transfection is likely due to aggregation of PROTAC with liposomal-based transfection reagent. Transfection efficiency data also showed that μDES electroporation resulted in enhanced loading efficiency, compared to the traditional cuvette-based electroporation technique (**Fig. 4A)**. The stability of loading and resulted PROTAC EVs were evaluated after EVs storage at 4°C for 24 hrs, at -80°C for 1 week, and at -80°C for 1 week following lyophilization. PROTAC EVs produced from μDES electroporation displayed long-term stability as the transfection efficiency was minimally altered by the various storage conditions and times (**Fig. 4B**). The EV particle size is slightly increased at the longer storage times at -80°C (**Fig. 4C**). However, EV particle concentration (**Fig. 4D**) was noticeably reduced, which is likely due to the EV aggregation. Therefore, freshly transfected EVs were utilized for all subsequent studies.

**Figure 4.**
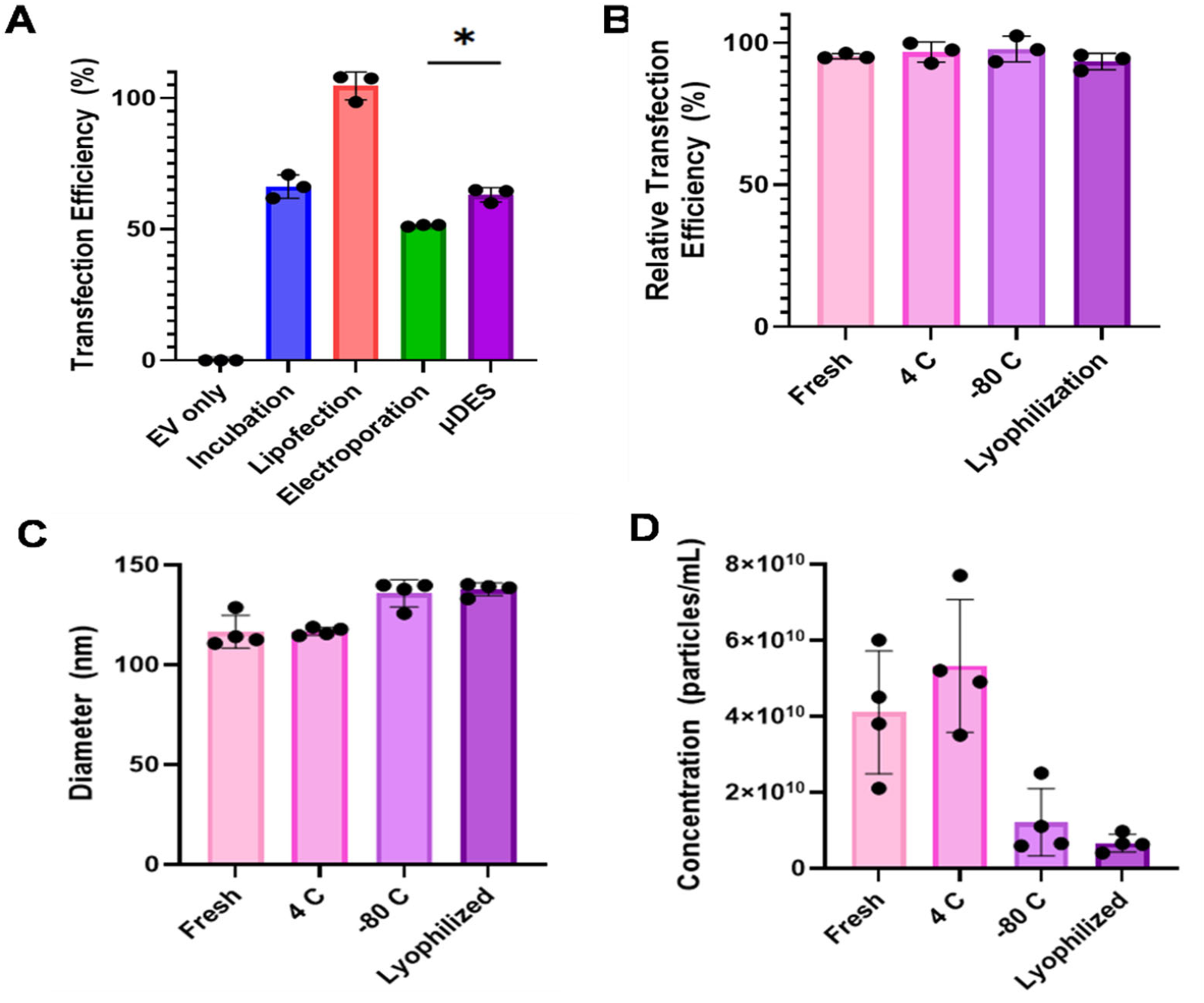
A) Transfection efficiency of PROTAC loaded into EVs. B) Transfection efficiency following storage at various conditions, representative of PROTAC loading stability. Effect of storage conditions on EV C) size and D) concentration.

### Assessment of HDAC protein degradation in vitro using transfected PROTAC EVs

In order to assess the therapeutic functionality of PROTAC loaded EVs in vitro, we compared the HDAC3/8 degradation capability using PROTAC EV transfected by various transfection techniques (**Fig. 5A**). Starting with the conventional loading methods of incubation, lipofection, and cuvette-based electroporation, EVs loaded with an initial input of 800 nM were compared to 800 nM of PROTAC alone for their ability to degrade HDAC3/8 in MDA-MB-231 cells (**Fig. 5B**). This concentration of YX968 is within the range shown in our previous studies to selectively and potently degrade HDAC3/8 in this cell line^33^. The unloaded EV control resulted in no protein degradation, while all other samples containing PROTAC, either alone or loaded in EVs, demonstrated significant protein degradation (**Fig. 5B**). The low concentration of 8 nM was also tested in parallel. In high concentration group, PROTAC loaded EVs through cuvette-based electroporation exhibited the strongest degradation of HDAC3 compared chemical transfection method using lipofection. Although low concentration is not effective for degradation on highly expressed HDAC3, electroporation loaded PROTAC EVs still exhibited the degradation ability on HDAC3 and 8 (**Fig. 5B**). In efforts to optimize the HDAC3/8 degradation of MDA-MB-231 cells treated with electroporated PROTAC EVs, further adjustments to EV and PROTAC concentrations were made in Figure 5C. Increasing the concentration of PROTAC used for EV loading from 16 nM to 100 nM led to a significant increase in HDAC3/8 degradation. Additionally, the highest concentration of EVs led to the highest level of HDAC8 degradation (**Fig. 5C**). These factors were then applied to assess the functionality of μDES produced PROTAC EVs. Similarly, increasing EV and drug concentration resulted in an increase in HDAC3/8 degradation (**Fig. 5C**). Therefore, the highest condition tested of 1.4×10^10^ EVs/mL and 100 nM PROTAC were utilized for following studies, which can achieve effective protein degradation in vivo and maintain reasonably attainable EV particle concentration. μDES produced PROTAC EVs displayed higher effective protein degradation in vitro, compared to conventional electroporation produced PROTAC EVs (Fig. 5C). Further validation was shown in Fig 5D, which proved μDES produced PROTAC EVs were responsive to HDAC 3 and 8 degradation along with cellular uptake time cycle. YX968-induced degradation is known to change over time^33^,The μDES produced PROTAC EVs were outperformed the other loading techniques. PROTAC loaded EVs demonstrated high potency in vitro, with degradative effects taking place as soon as 2 hrs following treatment to TNBC cells (**Fig. 5D**). Maximum HDAC3/8 degradation was observed within 10 hrs, with no signs of reversal of effects in the studied time periods. Therefore, PROTAC loaded EVs provide fast-acting and long-lasting protein degradation effects.

**Figure 5.**
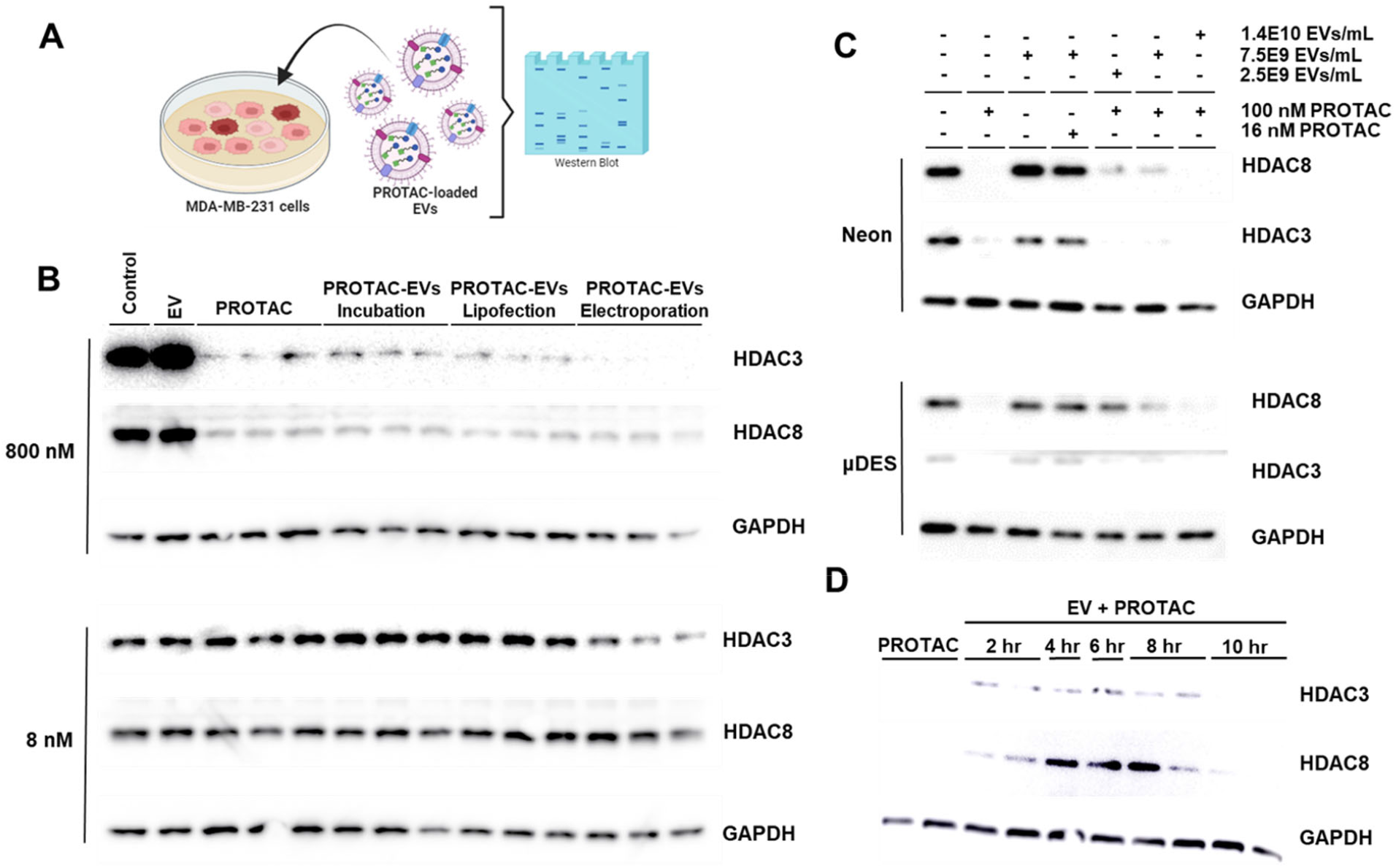
Functionality of PROTAC loaded EVs demonstrated through in-cell western blots of protein degradation. A) 800 nM and 8 nM PROTAC loaded in EVs following different conventional transfection methods. B) Various concentrations of PROTAC and EVs loaded through neon and μDES electroporation. C) Optimized μDES based PROTAC loaded EVs time dependent induction of protein degradation.

### Assessment of anti-tumor effect from μDES produced PROTAC EVs in TNBC mouse model

While the evaluation of protein degradation in vitro provides promising results in a controlled environment, there is often a disconnect between in vitro and in vivo results^34^. Therefore, in vivo assessments of PROTAC loaded EVs for degradation on HDAC3/8, and subsequently impart anti-tumor effects, were also conducted. Mice bearing MDA-MB-231 TNBC tumors were treated 3x per week with PROTAC loaded EVs over the span of almost 5 weeks (**Fig. 6A**). Mice were treated with unloaded EVs, PROTAC only, and PBS as controls. Measurements of the tumor volumes following the duration of the injections validated that unloaded EVs led to no change in tumor growth compared to the vehicle control, while PROTAC itself slightly lowered the final tumor volume (**Fig. 6B**). In contrast, the mice treated with PROTAC loaded EVs developed significantly smaller tumors with a reduced growth rate. Consequently, the PROTAC loaded EV treatment significantly extended the survival time of the mice (**Fig. 6C**). At the conclusion of the anti-tumor assessment, the tumors were collected, and protein degradation was analyzed (**Fig. 6D**). As expected, the tumors of mice treated with PROTAC loaded EVs had considerable HDAC3/8 degradation, while the tumors of mice treated with PROTAC alone did not show HDAC degradation. Therefore, EVs have the capability to enhance the therapeutic efficacy of PROTAC in vivo through augmented HDAC3/8 degradation and tumor inhibition.

**Figure 6.**
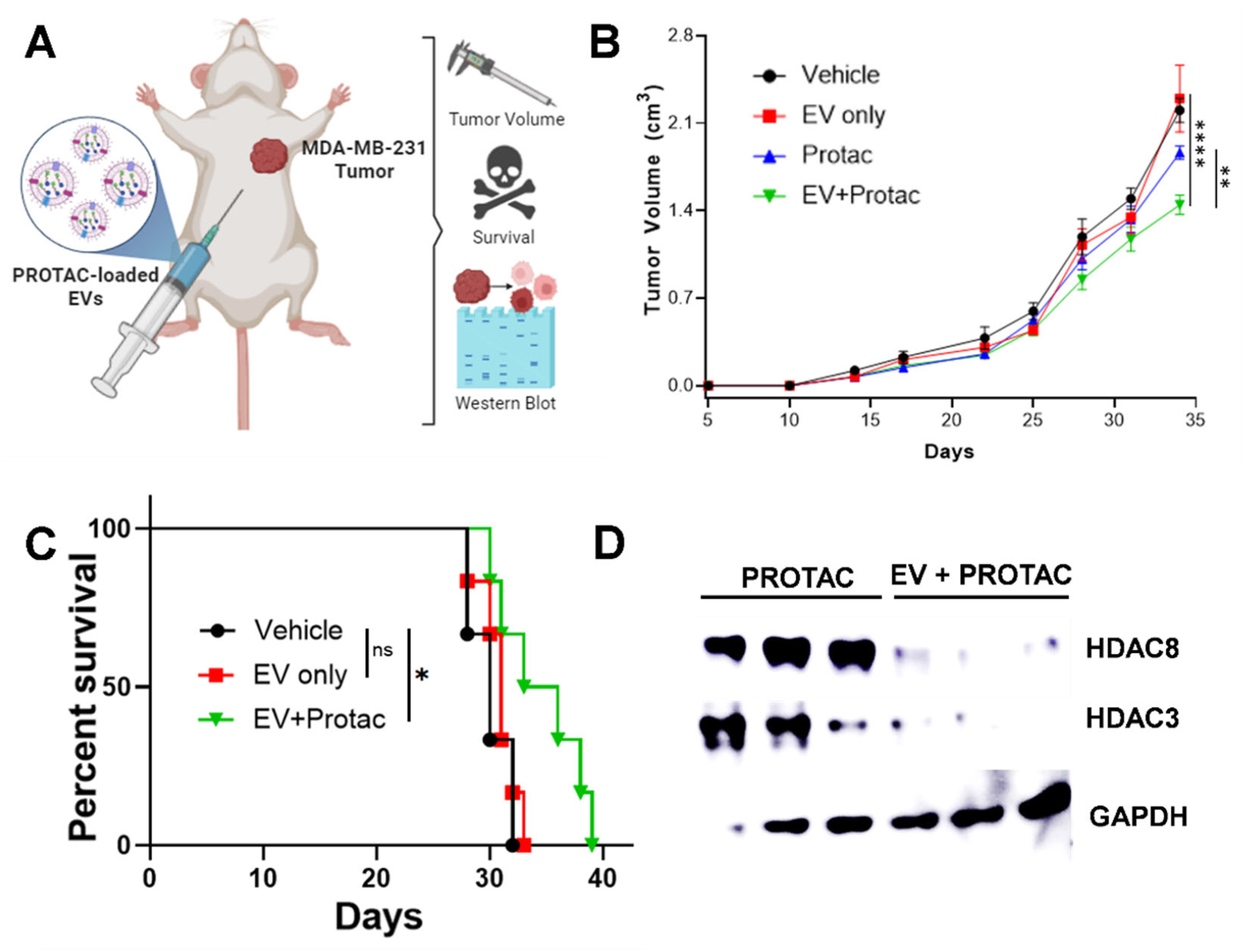
A) In vivo behaviors of μDES PROTAC loaded EVs in MDA-MB-231 breast cancer mouse models. Anti-tumor functionality of PROTAC loaded EV treatments as demonstrated through C) tumor size reduction, D) survival study, and E) protein degradation in the tumor.

### Assessment of safety profile in vivo from μDES produced PROTAC EVs

The potential toxicity of μDES produced PROTAC EVs were assessed to ensure safety following the treatment injections, with PROTAC and EV alone as the control group. To fully maximize the potential for identifying adverse effects, a high dose of both EVs and PROTAC were used (1.4×10^11^ EV/mL, 1000nM YX968). The concentration of both PROTAC and EV was increased by an order of magnitude over the previous studies to enhance exposure to the treatment associated adverse effects. The high doses were administered on day 0 and 7, body weights and complete blood counts were measured at 7 time points after the treatment and histological analysis was conducted on day 21 (**Fig. 7A**). No decline in body weights, regardless of treatment or gender, was observed (**Fig. 7B**). Additionally, there were no significant changes in hematological parameters, as white blood cells (**Fig. 7C**), lymphocytes (**Fig. 7D**), neutrophils (**Fig. 7E**), red blood cells (**Fig. 7F**), and platelets (**Fig. 7G**) remained relatively unchanged. Histological examination results using hematoxylin and eosin (H&E) staining demonstrated that the treatments did not induce any observable tissue damage to the analyzed organs (**Fig. 7H**). However, PROTAC drug only group displayed toxic effects from kidney and liver tissues. Based on these findings, EV loaded with PROTAC could reduce the PROTAC drug toxicity, which is able to improve the treatment safety.

**Figure 7.**
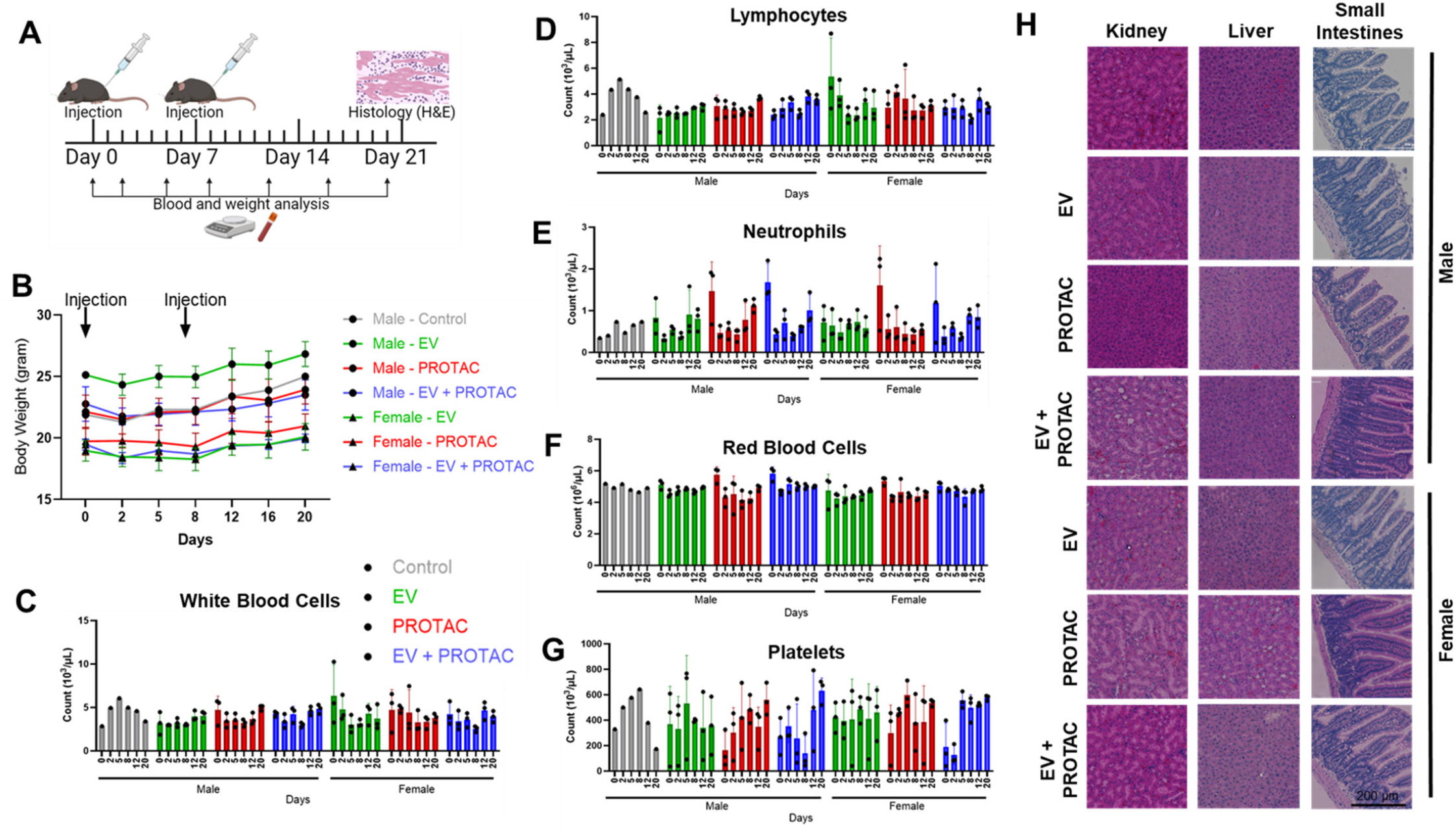
A) Schematic of toxicity assessment of high dose injections of μDES produced PROTAC EVs. B) Mice weight changes following treatment. Hematology analysis of blood cell components, including C) white blood cells, D) lymphocytes, E) neutrophils, F) red blood cells, and G) platelets. H) Impact of injection on tissue histology, including kidney, liver, and small intestine.

### Investigation of interspecies, multi-tumor biodistribution and protein degradation from μDES produced PROTAC EVs in TNBC multi-tumor model

EVs derived from various cell types have distinct characteristics ^35, 36^ that may impact EVs’ targeting behavior and the responses from target cells. The previously described experiments were conducted utilizing MDA-MB-231 cell derived EVs for PROTAC delivery to MDA-MB-231 cells. In order to better understand how the cell origin of EVs, and their targeting behavior and therapeutic efficacy, an alternative multi-tumor model was investigated. Human (MDA-MB-231) and mouse (PY8119) derived tumors were established in NOD-SCID interleukin-2 receptor gamma null (NSG) mice, which were then injected with a single dose of μDES produced PROTAC EVs derived from human cells (MDA-MB-231) (**Fig. 8A**). The mice were analyzed following 3 time points (6 hr, 15 hr, 24hr) to determine the HDAC protein degradation functionality in both tumor tissues. PROTAC alone and unloaded EVs were used as controls. The spleens of the mice did not display protein degradation for any of the applied treatments (**Fig. 8B**). Since tumors are the targeted tissue sites for PROTAC, this suggests that no off-targeted effects occur following the PROTAC EV treatment. In contrast, the human cell derived tumor exhibited fully targeted HDAC 3/8 protein degradation for 15 hr and 24 hr after treatments (**Fig. 8C**). However, the treatment of PROTAC alone did not show relevant targeted HDAC 3/8 protein degradation in any of those time frames. Interestingly, the mouse cell PY8119 established tumors did not undergo any protein degradation, regardless of the PROTAC treatment or time frames (**Fig. 8D**). This finding could be contributed to source-derived differences between the cellular origins of the EVs, PROTAC drug and tumor species. The biodistribution of μDES produced PROTAC EVs labeled with a near-infrared dye (DiR) was evaluated. The fluorescent signal of the EVs was captured immediately following injection and remained at strong levels up to 48 hrs post-injection (**Fig. 8E**). The PROTAC loaded EVs were relatively distributed in the same manner as the unloaded EVs, suggesting that the loading process does not impact the in vivo biodistribution behaviors of the EVs. To quantitatively monitor the fluorescence distribution, the tumors and major organs were harvested and imaged ex vivo (**Fig. 8F**). The EVs were primarily delivered to the spleen, liver, and both tumors. We suspect that degradation differences between tumor species may be involved with the origin of specific protein degradation. Nonetheless, the human cell derived PROTAC loaded EVs were demonstrated to be capable of distributing to MDA-MB-231 tumors, leading to efficient degradation of HDAC3/8.

**Figure 8.**
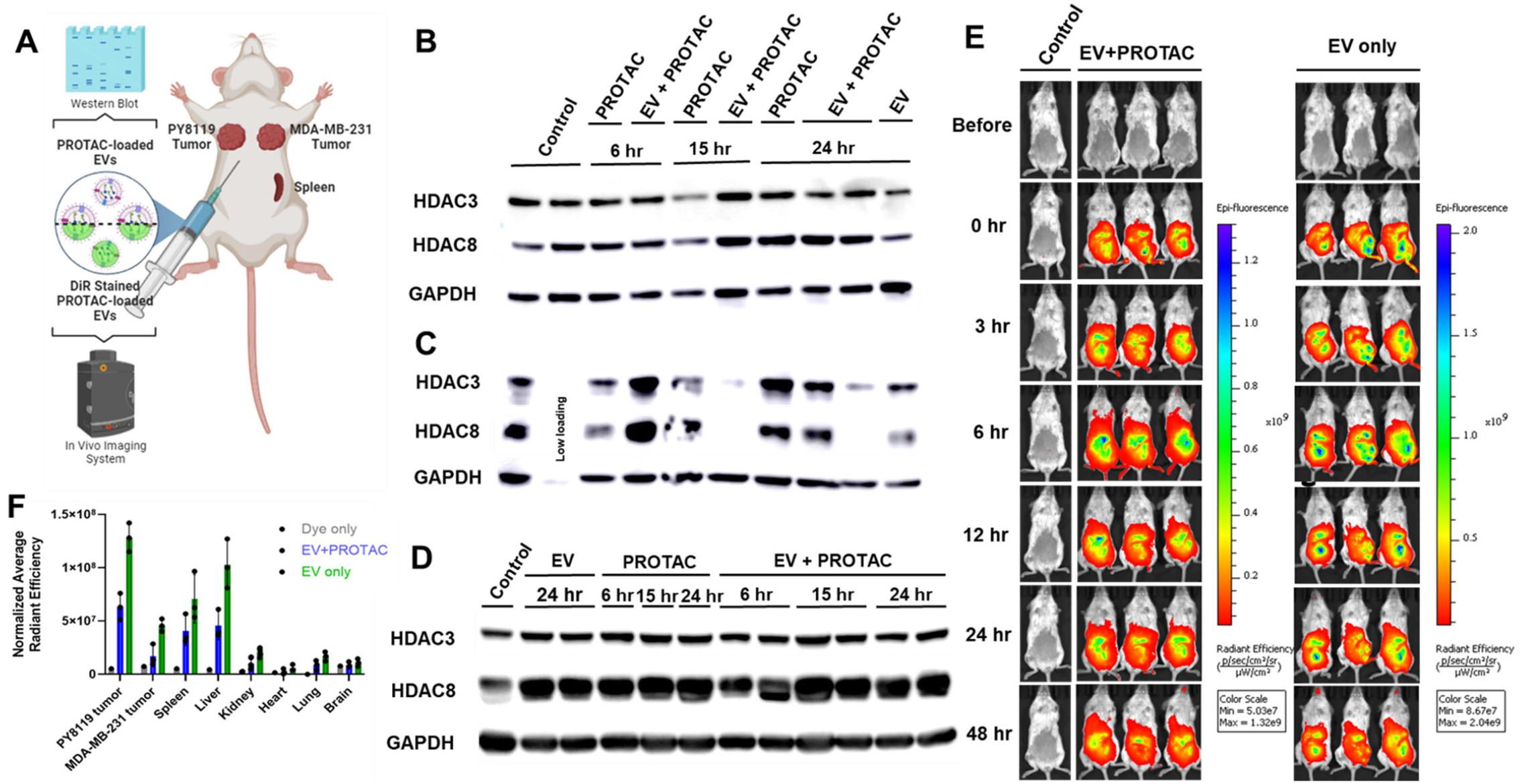
A) In vivo protein degradation and biodistribution behaviors of μDES based PROTAC loaded EVs in a multi-tumor model. Protein degradation assessed in the B) spleen, C) human MDA-MB-231 tumors and D) mouse PY8119 tumors. EV biodistribution in E) mice in vivo and F) organs ex vivo.

## Discussion

The clinical translation of PROTAC drug presented many challenges, such as low absorption, rapid drug degradation, and poor cell penetration in vivo. Thus, although YX968 was previously proven to effectively degrade HDAC3/8 in TNBC cell lines in vitro^33^, the adverse physicochemical and pharmacokinetic properties of the molecule hindered their effective anti-tumor function in vivo. Therefore, in efforts to improve the drug’s in vivo therapeutic function and achieve adequate HDAC3/8 degradation in vivo, in turn, leading to meaningful impacts on TNBC treatments, we introduced EV loaded PROTAC drug. Presently, EVs have been the emerging drug delivery vehicles which provide enhanced cargo protection and targeted delivery of carried cargos, allowing for enhancing therapeutic efficacy and anticancer therapy benefits. We investigated various transfection techniques for loading PROTAC into EVs, including incubation, chemical transfection through lipofection, and cuvette-based electroporation. In order to overcome the technology barrier for clinical translation, we also introduced the novel μDES transfection method for rapid, high throughput, efficient loading of PROTAC into EVs for the first time to achieve augmented protein degradation and effective anti-tumor function in TNBC mouse model.

Despite the high loading efficiency of PROTAC into EVs following lipofection (**Fig. 4A**), this transfection method greatly altered EV properties, especially surface charge and structural stability, which reduced EVs’ natural cell penetration ability and introduced significant cellular toxicity^37-39^ (**Fig. 5**). The electroporation techniques led to less drastic changes in retaining EV integrity (**Fig. 3**). In terms of the functionality of the PROTAC loaded EVs, the cuvette electroporation transfection method led to higher efficacy in degrading the HDAC3/8 as the proteins of interest, compared to lipofection and incubation (**Fig. 5B**). Similarly, the μDES and cuvette electroporation techniques both demonstrated degradation capability of HDAC3/8 in vitro (**Fig. 5C**), with improved protein degradation efficiency from μDES produced PROTAC EVs. On the other hand, continuous-flow microfluidic platform utilizing droplet-based EV electro-transfection, which can handle EVs in large throughput and high efficiency to enable clinical translation. Note that conventional cuvette processing volume is below ∼20 mL. With droplets generated in thousands per second, more than ∼20 mL per hr production rate can be achieved in a continuous-flow and automatic manner, which is ∼10^3^-fold enhancement on throughput over current transfection methods. Such efforts is critical for developing GMP compatibility and clinical grade therapeutic translation.

Freshly transfected EVs were prepared for each study, despite the fact that PROTAC loading into EVs remains stable for various storage conditions and time periods (**Fig. 4B**). Further, the optimum EV and PROTAC concentrations leading to the highest HDAC3/8 degradation were identified (**Fig. 5C**). More efforts will be pursued in the future to investigate the storage conditions and dosage formulation for improving shelf life as a drug.

We observed essential in vivo pharmacokinetic behavior of PROTAC EVs compared to PROTAC drug only. The in vivo treatments of PROTAC alone exhibited only minimal degradation of HDAC 3/8, and such minimal degradation was reversed after 24 hrs (**Fig. 6D/8C**). In contrast, in vivo treatments of PROTAC EVs displayed complete degradation of HDAC3/8 at 15 hr and 24 hr, with no reversal of degradation effects in the time frame studied. These in vivo HDAC3/8 degradations were only identified in the tumors, with no effects detected to the spleen (**Fig. 8B-D**). We did not observe the HDAC3/8 degradation from a non-homologous cell source to the EVs (**Fig. 8B-D**). Although, the observed EV biodistribution behavior showcased that EVs travel in large quantities to each of these tissues (spleen, human MDA-MB-231 tumor, and mouse PY8119 tumor) (**Fig. 8E/8F**). However, biodistribution to a tissue does not indicate that EV uptake occurs, which is essential for PROTAC delivery and subsequent protein degradation. The results may suggest that the EVs promote specific uptake for release of the carried payload. Further investigation will be conducted to understand this observation.

Since PROTAC loaded EVs lead to effective HDAC3/8 degradation in vivo, and HDAC3/8 play important roles in tumorigenesis and cancer progression, TNBC tumor growth inhibition following PROTAC loaded EV treatment can also be enhanced. This was demonstrated as injections of PROTAC alone only led to minimal effects on tumor volume, in contrast, PROTAC loaded EVs were capable of significantly reducing tumor volume (**Fig. 6B**). Consequently, the mice treated with PROTAC loaded EVs had prolonged median survival times by 4.5 days (**Fig. 6C**). The improved anti-tumor efficacy of the PROTAC loaded EVs are likely a result of the enhanced PROTAC delivery to the tumor, enabling the improved HDAC3/8 degradation. Additionally, the intended, desired anti-tumor effects were not accompanied by any adverse effects compared to PROTAC alone group (**Fig. 7**). The safety and efficacy demonstrated by the PROTAC loaded EVs provides important insight into EVs’ potential clinical application as promising anti-cancer drug delivery vehicles.

The results from this study have proven that EVs, as drug delivery systems, can improve the therapeutic function of PROTAC in mouse model, allowing them to achieve much improved efficacy and safety in vivo. As reported, ∼90% of drug developments fail in clinical studies, which could be attributed to an imbalance of drug dose, efficacy, and safety, as well as a biological discrepancy between in vitro and in vivo models^40^. Therefore, our study could provide a viable solution to enhance clinical drug development success rates. Drug delivery through EVs could be implemented to ensure increased targeted delivery and therapeutic performance of the drug. The enhanced in vivo effects of PROTAC following loading into EVs has promising implications for applying to other drugs as well. Other therapeutics with advantageous characteristics, but unfavorable behaviors in vivo, can have a revived opportunity for clinical approval through the enhanced functionality granted by EV delivery. Therefore, the μDES platform for engineering drug loaded EVs is a novel approach for the delivery of therapeutics and may be a promising strategy for future EV applications to deliver the clinical success of drugs.

## Methods

### Cell culture

MDA-MB-231 human breast cancer cells, as well as PY8119 murine breast cancer cells, (ATCC) were cultured in Dulbecco’s Modified Eagle Medium (DMEM, Gibco) containing 10% exosome-depleted fetal bovine serum (Gibco) and 1% penicillin-streptomycin (Sigma). Cells were passaged when confluency reached approximately 80% by removing culture media, detaching cells using 0.25% Trypsin-EDTA (Gibco), pelleting cells through centrifugation at 300 x g for 5 min at 22 C, and resuspending in fresh media at a subcultivation ratio of 1:3. Cultures were maintained in a humidified incubator with 5% CO2 supply at 37 °C and all studies were performed during the logarithmic phase of cell growth.

### EV purification and characterization

Cell culture media was collected 48 hrs following subculture for subsequent EV isolation through differential centrifugation. First, cells were removed from the media by centrifugation at 300 x g for 10 min at 4 C. The supernatant was then centrifuged at 2000 x g for 30 min at 4 C to remove cell debris. Apoptotic bodies were then removed through centrifugation at 10,000 x g for 30 min at 4 C. The media was then transferred to thick wall ultracentrifuge tubes (Thermo Scientific) and EVs were collected by ultracentrifugation at 100,000 x g for 90 min at 4 C (Sorvall MTX150 Micro-Ultracentrifuge, Thermo Scientific). The EVs were washed with 8 mL cold Dulbecco’s phosphate-buffered saline (PBS, Sigma) and isolated again through ultracentrifugation at 100,000 x g for 90 min at 4 C. The EV pellet was resuspended in PBS for analysis and further experimentation.

Isolated EV size, zeta potential, and particle number was analyzed by nanoparticle tracking analysis using the ZetaView Quatt Nanoparticle Tracking Analyzer – PMX420 instrument (Particle Matrix). Machine calibration was performed using 100 nm polystyrene beads diluted 1:250,000. The EV solution was diluted 1:1,000, a total of 1 mL was injected into the measuring cell, and 11 cell positions with 5 cycles per position were measured. The zeta potential was measured in pulsed mode using 2 positions, with 2 cycles per position. The capture settings were: sensitivity 80, frame rate 60, shutter 163, minimum trace length 15. A washing step using PBS was conducted between each EV measurement. Analysis was then performed with ZetaView software 8.05.17 SP7. The morphology of the EVs was determined by transmission electron microscopy (FEI Spirit TEM 120 kV) imaging. Briefly, ultrathin copper grids coated with 400 mesh carbon film (Electron Microscopy Science) were treated with glow discharge. Then, 15 μL of the isolated EV solution was added onto the grids for 5 min. The grids were then washed with distilled water and negatively stained with filtered 2% aqueous uranyl acetate for 2 min.

### µDES device fabrication

The design for the µDES device master mold was uploaded into the Utility software (version 6.3.0.t3) of the MiiCraft Ultra 50 3D Printer (Creative CADworks). The master mold was printed in master mold resin (CADworks3D) and cleaned in absolute ethanol, 200 proof (fisher scientific) overnight. The mold was then cured for 3 min on each side using the Professional CureZone (Creative CADworks) before casting with polydimethylsiloxane (PDMS) using the SYLGARD 184 Silicone Elastomer Kit (Dow). Air bubbles were removed by incubating the device in a desiccator cabinet (SP Bel-Art) connected to a vacuum pump (WOB-L 2534B-01, Welch) for 1 hour. The device was then heat cured in a 75 C Heratherm oven (Thermo Scientific) for 3 hours. Following device removal and cooling, inlets and outlets were formed with a 1 mm biopsy punch (Miltex). The device was then bound to a glass slide (fisher scientific) following surface cleaning and activation for 10 sec using a plasma cleaner (Harrick Plasma) connected to a vacuum pump (DVP).

Polytetrafluoroethylene tubing (Cole-Parmer) was inserted into the inlets and outlets. Then, platinum iridium wire (Surepure Chemetals) connected to 0.644 mm solid electrical wire (Tuofeng) was added into the electrode channels. All surfaces were then sealed with PDMS to prevent leaking.

### EV transfection

Chemical transfection was performed using the Lipofectamine 3000 Transfection Reagent (Invitrogen). Briefly, 100 nM YX968 (2.19 µg) was combined with 2 µL/µg P3000 Enhancer Reagent and Opti-MEM media (gibco). Then, Lipfectamine 3000 Reagent, at a volume 3 times YX968 (6.57 µL) was diluted in Opti-MEM and added to the previous solution.

Following 10 min incubation at RT, the mixture was added to isolated EVs diluted to 1.4×10^10^ EVs/mL for an additional 24 hrs of incubation.

Direct physical transfection was performed by simply combining 100 nM YX968 with isolated EVs diluted to 1.4×10^10^ EVs/mL and incubating for 1 hr.

Traditional cuvette-based electroporation was conducted using the Neon electroporation device (Invitrogen). EVs were diluted to 1.4×10^10^ EVs/mL in Cytoporation Low Conductivity Media T (CytoT, BTXpress) and combined with 100 nM YX968 and 50 nM trehalose (Acros Organics). The EV media was loaded into 100 µL reaction tips and inserted into transfection tubes containing 3 mL of 10% 111.9 mS/cm Conductivity Standard (Thermo Scientific) in water with 50 nM trehalose. The EV solution was pulsed once at 1500 V for 20 millisec and then allowed to rest at RT for 30 min.

µDES-based electroporation was performed by connecting the µDES device to the iFlow Touch Microfluidic Pump System (PreciGenome) for pressure-controlled flow and to the DC Power Supply (GW Instek) for electroporation. First, the device channels were washed by flowing through absolute ethanol and stabilized by flowing through the 10% conductivity standard. Then, buffer in oil droplets were formed by connecting EVs diluted to 1.4×10^10^ EVs/mL in CytoT buffer and 50 nM trehalose to the µDES device along with 2 wt% 008-FluoroSurfactant in FC40 oil (Ran Biotechnologies). The droplets are electroporated at 30 V and allowed to rest at RT for 30 min. The size and morphology of the droplets were characterized using the Cytation 5 imager (BioTek). Then, to break the emulsion and remove the oil phase to collect EVs, the solution was centrifuged at 2000 x g for 5 min at 4 C.

### Purification of loaded EVs

EVs were washed through ultrafiltration to remove unincorporated YX968. In short, the EVs were centrifuged at 10,000 x g for 30 min at 4 C in Amicon Ultra-0.5 Centrifugal Filter Units with a molecular weight cut off of 10 kDa (sigma). Then, fresh ultrapure distilled water (Invitrogen) was added to the EV solution which were centrifuged again. The washing process was repeated 3 times. The remaining EVs in the filter were collected and resuspended.

### Loading efficiency quantification

To assess the concentration of YX968 loaded into EVs, we utilized liquid chromatography with tandem mass spectrometry (LC-MS/MS) for quantification. For EV lysis and protein denaturation, we combined 60 µL of the EV solution with 50 µL of methanol, and 100 µL of acetonitrile. Then, 6 µL of the internal standard at 250 ng/mL was added for further sample preparation. Following brief sonication and centrifugation, 150 µL of the supernatant was transferred for analysis with the 3200 QTRAP triple quad/linear ion trap LC-MS/MS (Applied Biosystems).

For the assessment of loading efficiency over time, the EV solution with 50 nM trehalose was stored at 4 C for 24 hrs and at -80 C for 1 week, both in solution and following lyophilization. Following storage, the EVs underwent ultrafiltration using a 10 kDa molecular weight cut off for the removal of leached YX968. Then EV lysis and sample preparation for LC/MS/MS was performed as described above.

### Western blot analysis

For the in vitro western blot analyses, MDA-MB-231 cells were seeded in a 48-well plate (50,000 cells/well), allowed to grow for 24 hrs, and treated with the EV or YX968 solutions as indicated in relevant figures. 8 hrs after treatment, or as stated elsewhere, the media was removed, and the cells were lysed by adding RIPA buffer (80 µL, Thermo Scientific) and shaking at 500 rpm for 1 hr. The cell lysates were then mixed with 6x SDS samples buffer (16 µL, Bio-Rad), heated to 95 C for 5 min, and cooled to RT. After brief centrifugation, 15 µL of the lysates were then loaded on a 4-20% gradient gel (Novex Tris-Glycine Mini Gels, ThermoFisher) and run at 200 V for 45 min at RT. The proteins were electrotransferred to a polyvinylidene fluoride membrane (sigma) using an Trans-Blot Turbo Transfer System (Bio-Rad). The membranes were blocked with 5% nonfat milk, incubated with primary and secondary antibodies, and washed with a tris-buffered saline solution containing 0.1% tween-20. The primary antibodies used include anti-HDAC3 (Abcam, 1:10,000 dilution), anti-HDAC8 (ProteinTech, 1:5,000 dilution), and anti-GAPDH (ProteinTech, 1:10,000 dilution) diluted in 1% nonfat milk and 0.05% sodium azide (sigma). The secondary antibodies included HRP-linked anti-mouse and anti-rabbit IgG antibody (1:10,000 dilution, Cell Signaling Technology). Finally, the proteins were detected by the Amersham Imager 680 (GE) using a Chemiluminescent HRP Substrate detection kit (sigma).

For the in vivo western blot analyses, murine tissue samples were homogenized through cryogenic grinding using a pestle and mortar containing liquid nitrogen over dry ice. The tissue homogenates were suspended in RIPA Lysis and Extraction Buffer (500 μL, Thermo Scientific) supplemented with 10 uL/mL of Halt Protease and Phosphatase Inhibitor Cocktail 100x (Thermo Scientific) and briefly ultrasonicated using a dismembrator homogenizer with probe (Fisher Scientific) at 40 kHz in 5-10 sec increments. The lysates were then centrifuged at 15,000 rpm at 4 C for 5 min and the supernatant was added to SDS buffer, following the western blot steps as described above.

### Animal studies

All procedures were approved by the Institutional Animal Care and Use Committee (IACUC) of the University of Florida and followed the guidelines established by NIH. 7–9-week-old C57BL/6J and NOD-SCID interleukin-2 receptor gamma null (NSG) mice were purchased from Jackson Laboratories. For tumor models, 1×10^6^ MDA-MB-231 cells, and 5×10^5^ PY8119 cells, were resuspended in PBS and implanted into mice subcutaneously. Once tumors reached 1 cm in diameter (about 10 days), mice were treated with the relevant injections.

For in vivo biodistribution studies, YX968-loaded EVs were incubated with 1,1’-Dioctadecyl-3,3,3’,3’-Tetramethylindotricarbocyanine Iodide (DiR, Invitrogen) at a final staining concentration of 5 μg/mL for 20 min at RT for membrane labeling. The unincorporated dye was removed through ultracentrifugation at 100,000 x g for 90 min at 4 C. EVs were resuspended in PBS to 1.4×10^10^ EVs/mL and filtered with a 0.2 µm syringe filter (Corning). As a control, unloaded EVs and PBS without EVs were subjected to the same DiR treatment. NSG mice, with pre-established MDA-MB-231 and PY8119 breast cancer tumors, were injected intraperitoneally (IP) with the samples in 0.1 mL PBS. To evaluate biodistribution, the mice were imaged 0, 3, 6, 12, 24 and 48 hrs following treatment injection using IVIS Spectrum Imaging System (Xenogen, Caliper Life Science). 48 hrs post treatment injection, the mice were euthanized and the heart, liver, lung, kidney, spleen, brain, and tumors were harvested for ex vivo imaging. The imaging parameters include: excitation = 745 nm, emission = 800 nm, field of view = 13.5 cm, fluency rate = 2 mW cm^-2^. The camera was set to a maximum gain, a binning factor of 8, F stop = 1, and an exposure time of 1 s. The data was analyzed with the IVIS software (Living Image Software for IVIS).

For in vivo tumor inhibition studies, YX968-loaded EVs were suspended in PBS to 1.4×10^10^ EVs/mL and filtered with a 0.2 µm syringe filter. NSG mice with pre-established MDA-MB-231 tumors were treated with the EVs by intraperitoneal injection 3x /week. Unloaded EVs, 100 nM PROTAC, and PBS were used as controls. Tumor size and body weights were measured 2-3x / week.

For the in vivo toxicity assessment, non-tumor-bearing C57BL/6J mice were injected with high dose treatments of 1.4×10^11^ EV/mL YX968-loaded EVs, prepared using a starting concentration of 1000 nM YX968, on days 0 and 7. Unloaded EVs and 1000 nM PROTAC were used as controls. Blood samples were taken from the submandibular (facial) vein before treatment and 2-3x /week for hematological analysis. Blood samples were prepared in PBS with 10 mM EDTA and blood indices were analyzed using an automated hematology analyzer (Element HT5; Heska). Mice body weights were also measured 2-3x /week. Two weeks following the final treatment, mice were euthanized and tissues were collected for histological analysis. Tissues were fixed in 10% formalin (SF98-4; Thermo Fisher Scientific), hydrated through xylene and a series of ethanol, and stained with hematoxylin and eosin (H&E) on slides.

### Statistical Analysis

Statistical analysis was conducted using the Student’s two-tailed t-test and one-way analysis of variance (ANOVA). P values of 0.05 or less were considered statistically significant. Statistical analysis was performed using JMP Pro 16.1.0. Data is expressed as means ± standard deviations.

## Acknowledgments

The authors would like to thank the Interdisciplinary Center for Biotechnology Research (ICBR) core facility at the University of Florida. We acknowledge funding support from NIH 1R35GM133794, and NIH TARGETED Challenge to Dr Mei He.

